# Differential association of SARS-CoV-2 IgG responses with anti-OC43 IgG in a Senegalese cohort

**DOI:** 10.1101/2025.04.07.647508

**Authors:** Rokhaya Faye, Adji Astou Mbow, Billo Tall, Aboubacry Gaye, Amadou Moustapha Ndoye, Rokhaya Gueye, Dieynaba Diallo, Francine Ndebane Faye, Cheikh Talla, Ndongo Dia, Bababcar Mbengue, Stéphane Pelleau, Makhtar Niang, Moussa Seydi, Amadou A. Sall, Fabien Taïeb, Cheikh Loucoubar, Michaël White, Thomas Poiret, Inés Vigan-Womas

**Affiliations:** Institut Pasteur de Dakar, Immunophysiopathology and Infectious Diseases Department, Dakar, Senegal; Institut Pasteur de Dakar, Epidemiology, Clinical Research and Data Sciences Department, Dakar, Senegal; Institut Pasteur de Dakar, Virology Department, Dakar, Senegal; Université Cheikh Anta Diop de Dakar, Service d’Immunologie FMPO, Dakar, Senegal; Institut Pasteur, Infectious Disease Epidemiology and Analytics G5 Unit, Paris, France; Department of Infectious and Tropical diseases, Fann University Hospital, Dakar, Senegal; Karolinska Institute, Department of Medicine Huddinge (MedH), Stockholm, Sweden; Institut Pasteur, Paris, France

**Author notes:** Corresponding author: /.

**Keywords:** SARS-CoV-2, COVID-19, Human coronaviruses (HCoVs), OC43, IgG antibody kinetics, Cross-reactivity

## Abstract

Numerous studies elucidated the kinetics of the humoral immune response post-SARS-CoV-2 infection. However, in sub-Saharan Africa, the evolution of SARS-CoV-2 IgG antibody responses and their interaction with pre-existing seasonal human coronavirus (HCoVs: OC43, 229E, NL63, HKU1) immunity remain underexplored.

A prospective cohort study was conducted in Senegal during the first year of the COVID-19 pandemic (March to December 2020). A total of 204 patients with laboratory-confirmed COVID-19 were included. Patients were classified as symptomatic (n=157) or asymptomatic (n=47) based on clinical presentation. Plasma samples (n=705) were collected over 6 months from SARS-CoV-2 positive individuals. IgG levels against SARS-CoV-2 and HCoVs were measured using a multiplex bead-based assay.

Among the 204 participants included (95 [46.6%] female, median age, 44 [7–95]), SARS-CoV-2 IgG were detectable 6 months post-infection, peaking at 1 month for most antigens, except for Spike (S), which peaked at 3 months. Elderly patients (>60 years) exhibited higher IgG levels against both SARS-CoV-2 and HCoVs. Symptomatic patients had higher IgG levels than asymptomatic individuals, especially for WTS, RBD, S2, and N. Anti-HCoV IgG levels remained stable post-infection, with OC43 peaking at week 3 in symptomatic patients. A positive correlation was found between anti-SARS-CoV- 2 and anti-OC43 IgG in symptomatic patients.

The study highlights persistent SARS-CoV-2 IgG antibodies for up to 6 months and suggests a link between pre-existing HCoV-OC43 immunity and COVID-19 outcomes in Senegal. These findings could help shape future vaccine strategies, considering the influence of circulating HCoVs on long-term protection against SARS-CoV-2.

**Author summary:** Understanding how our immune system responds to SARS-CoV-2, the virus responsible for COVID- 19, is essential for guiding public health countermeasures and informing vaccine development strategies. In our study, we monitored, in COVID-19 patients, the evolution of IgG antibody responses against SARS-CoV-2 structural proteins over a six-month period. Additionally, we examined how previous exposure to common seasonal coronaviruses might influence immune responses to SARS-CoV-2. Conducting this research in an African context is particularly important, as data on immune responses to SARS-CoV-2 in this region are scarce.

Our results provide valuable insights into the complex interplay between immune responses elicited by SARS-CoV-2 and pre-existing immunity from seasonal circulating coronaviruses. These findings enhance our understanding of immune memory and cross-reactivity, two critical factors for assessing long-term protection and optimizing vaccine strategies. By shedding light on the dynamics of antibody responses over time within a sub-Saharan population, our research contributes to the global effort aimed at developing effective interventions against COVID-19 and preparing for future coronavirus outbreaks.

## Introduction

Although several years have passed since the emergence of COVID-19 [1], our understanding of the immune response to SARS-CoV-2 remains incomplete; particularly regarding the role of IgG antibodies in long-term immunity and their interaction with other human seasonal coronaviruses (HCoVs). Characterization of the nuances of immune responses to SARS-CoV-2 is critical for optimizing public health strategies and development of updated vaccines.

Dynamics of antibody (IgG) responses following SARS-CoV-2 infection typically peak shortly after infection and can persist for months, albeit with gradual declines over time [2–4]. The IgG kinetics are known to vary with more severe cases often generating higher and more sustained antibody levels compared to mild or asymptomatic cases [5–8]. However, few studies have been conducted in low- and middle-income countries, particularly in Sub-Saharan Africa [9, 10].

In addition to the highly pathogenic SARS-CoV-2 coronavirus, four seasonal human coronaviruses (HCoVs) are known to cause mild respiratory infections: alphacoronaviruses (229E and NL63) and betacoronaviruses (OC43 and HKU-1), to which SARS-CoV-2 also belongs [11, 12]. The cross-reactivity between antibodies generated against SARS-CoV-2 and these seasonal HCoVs may determine how pre-existing immunity can influence the severity and outcomes of COVID-19 infections: While some studies suggested that prior exposure to HCoVs may influence the course of COVID-19 by either providing partial protection or modifying immune responses [13, 14], others described minimal cross-reactivity and no significant effect on SARS-CoV-2 immunity [15]. The variability in findings across studies may stem from differences in populations, methodologies, and settings.

HCoVs have seasonally circulated in Senegal, underscoring their potential implications for public health [16, 17] allowing to investigate the potential role of pre-existing HCoV immunity in influencing SARS-CoV-2 infection outcomes. Considering the heterogeneity of antibody responses in different populations and the limited data from Africa, with this study we provide the first in-depth longitudinal analysis of SARS-CoV-2 IgG responses and their potential cross-reactivity with the four seasonal HCoVs over a six-month period in a West African population. The kinetic of the SARS-CoV-2 humoral response was analysed through a multidisciplinary approach using multiplex serological assays, epidemiological data, and clinical records, to contribute to a more comprehensive understanding of the immune response to SARS-CoV-2 in an African context.

## Results

### Patient characteristics

This study analyzed humoral immune responses in a cohort of 324 individuals that included 204 patients infected by SARS-CoV-2, and a pre-SARS-CoV-2 (pre-pandemic) control cohort composed of 120 individuals from Dielmo village sampled in July 2018. The epidemiological and clinical characteristics of the SARS-CoV-2 cohort are described in **Fig 1B** and in **Table 1 in S1 Text**. For patients infected by SARS-CoV-2, the median age of this study population was 44 years ranging from 7 to 95 years and was divided by quartiles for analysis. Most patients were male (53.4%) and symptomatic (77%). There was a significant association of symptoms occurrence with age groups (**χ**^2^=9.116, p=0.027) but not with gender (**χ**^2^=0.496, p>0.05) (**Fig 1B**).

**Fig 1.**
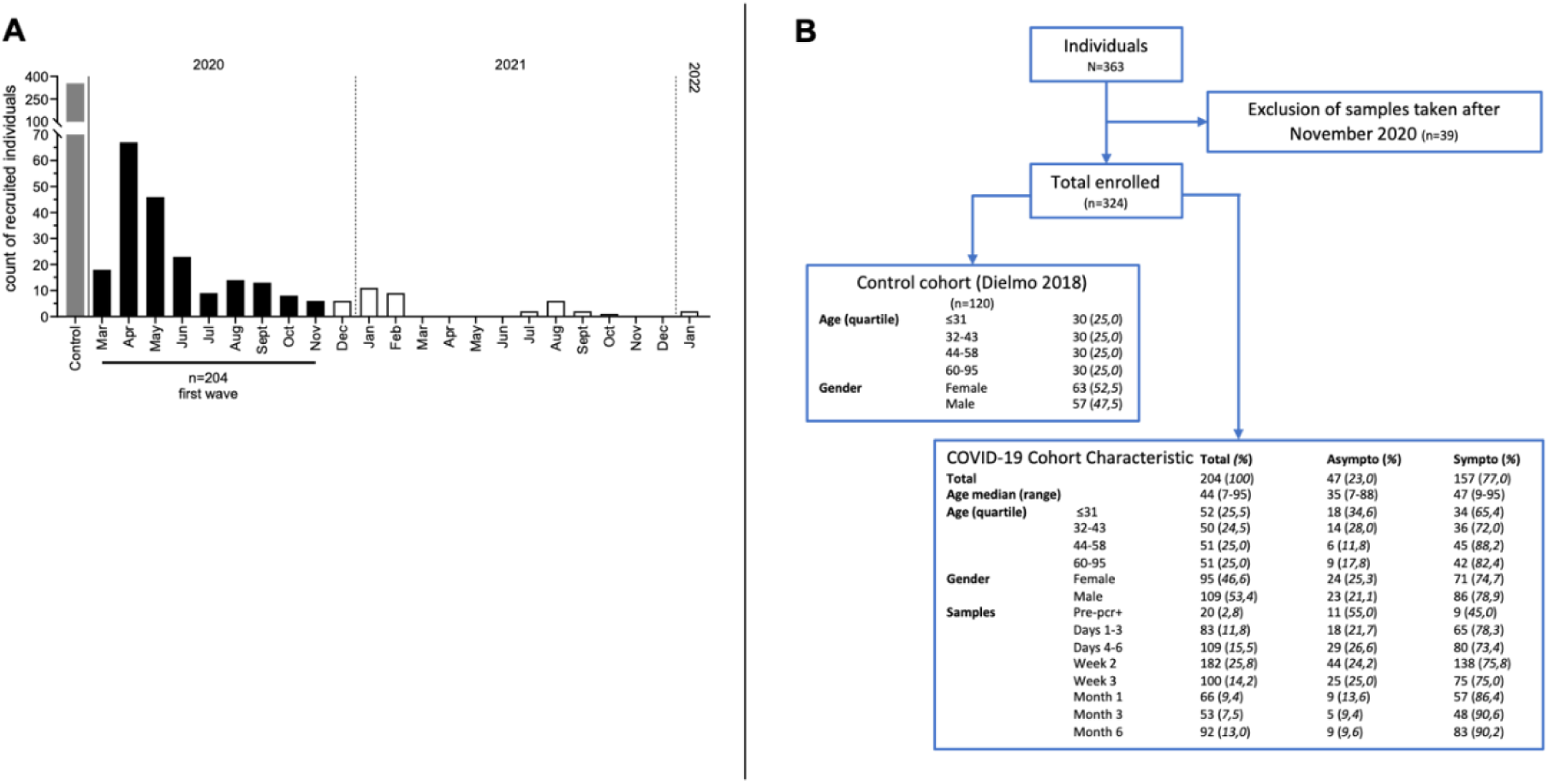
**Study overview and patients’ cohort information**. ***A***, Detailed cohort inclusion of post-COVID-19 first wave in Senegal between March and November 2020. A pre-SARS-CoV-2 (pre-pandemic) control cohort composed of individuals from Dielmo village sampled in July 2018 was also included. ***B,*** Inclusion/exclusion flowchart of control and patient samples in the study with patients and sampling information.

For antibody kinetics analysis, a total of 8 sampling timepoints from the 1^st^ positive PCR test were taken: pre-1^st^ PCR+, day 1-3, day 4-6, week 2, week 3, months 1, 3 and 6 post-1^st^ PCR+ samples. Overall, 705 samples were included in this longitudinal analysis. The number of samples analyzed by timepoints varied from 20 to 182 per timepoints with the highest proportion (25.8%) collected at week 2 post-1^st^ PCR+. A significant relationship was found between the timing of sample collection and the disease status of the patients (χ2=31.17, p<0.001): a smaller number of samples from asymptomatic patients was collected from month 1 and later post-1st PCR+ (representing <13,6% of samples, **Fig 1B**).

### Overall kinetics of the SARS-CoV-2 and seasonal HCoV IgG-responses post-infection

We monitored the dynamic of anti-SARS-CoV-2 and anti-HCoV IgG antibody levels from the 1^st^ PCR+ test and up to 6 months. The assay specificity was validated by hierarchical clustering of IgG levels from 11 asymptomatic patients and 9 symptomatic patients pre-1^st^ PCR+ samples (**S1 Fig**) showing no clustering based on the patients’ symptomatic status. From these pre-PCR+ reference samples, the proportional changes of anti-SARS-CoV-2 and anti-HCoV IgG antibody levels showed a significant rapid increased response post-1^st^ PCR+ against SARS-CoV-2 antigens (> 10-fold minimum at 1 month post-1^st^ PCR+) but not against HCoV antigens (**S2 Fig**).

To assess the kinetics and durability of the anti-SARS-CoV-2 antibody response, we analyzed a total of 1,183 samples from 682 individuals, including 705 samples collected from SARS-CoV-2 PCR+ patients and 120 pre-pandemic samples (**Fig 2**). IgG concentrations against SARS-CoV-2 proteins were significantly higher in patients’ cohort as compared to the age-sex matched pre-pandemic control cohort. Overall, from pre-1^st^ PCR+ timepoint, a significant increase of anti-SARS-CoV-2 IgG plasma levels were observed and remained detectable up to 6 months. Apart from WTS, which peaked at month 3 post-1^st^ PCR+ (**Fig 2A**), the peak levels of specific IgG were found at month 1 post-1^st^ PCR+ against all SARS-CoV-2 proteins (**Fig 2A**). From the peak IgG level, anti-N and anti-ME responses exhibited a significant decrease between 1- and 6- months post-1^st^ PCR+ (**Fig 2A**). Minimal variation in the kinetics of anti-HCoV IgG levels was found between pre-pandemic and SARS-CoV-2 samples (**Fig 2B**). Only IgG levels against OC43 Spike protein showed significant changes with a peak in antibody level at month 1 post-PCR+ (**Fig 2B**, p<0.05).

**Fig 2.**
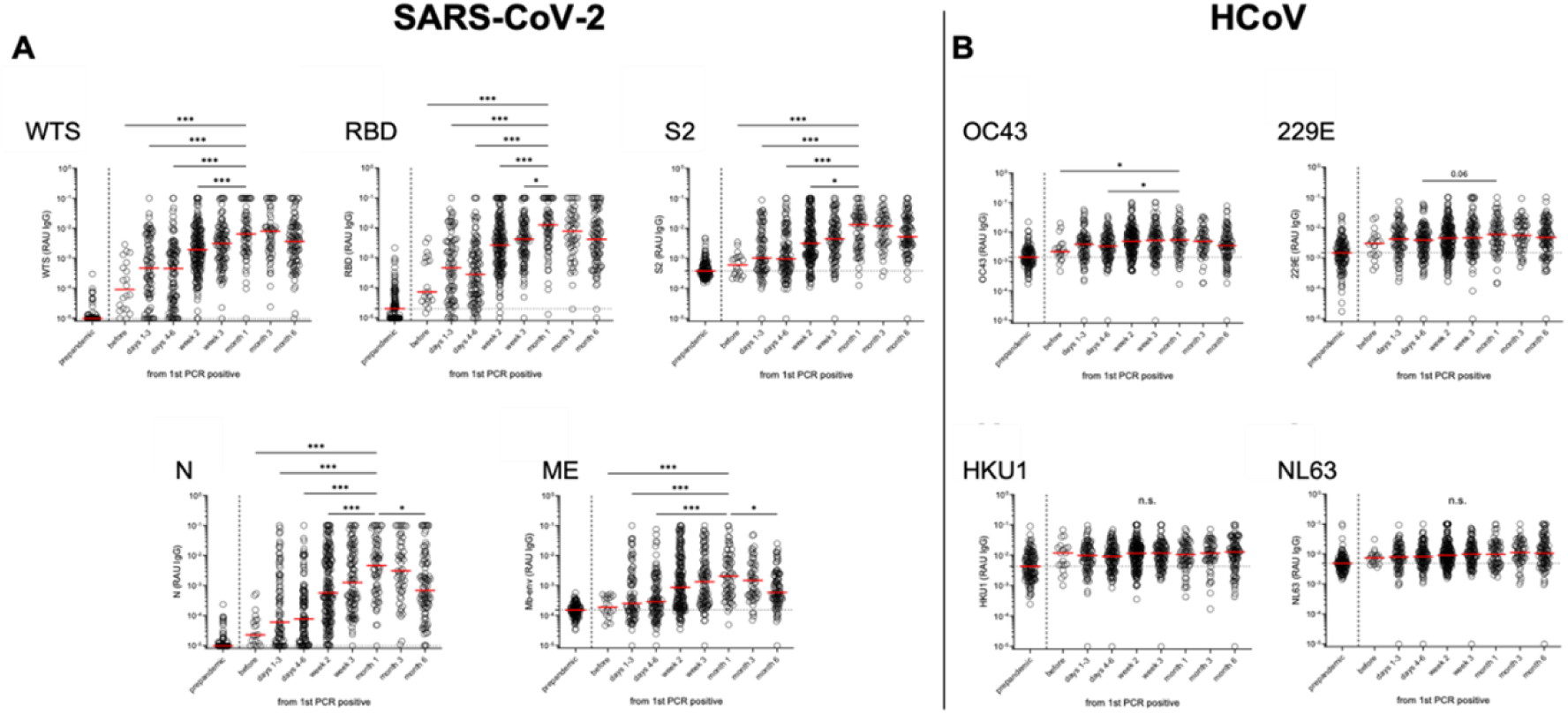
Six-month kinetics of SARS-CoV-2 IgG level. Individuals (n=204) IgG RAU kinetic responses to 5 SARS-CoV-2 (***A***) and 4 HCoV (***B***) different protein antigens. Kruskal-Wallis test with Dunn’s multiple comparison was used to compare values of group of individuals overtime. Statistical analyses were performed using the peak of the response as reference, i.e. at month 1 post-PCR+ except WTS antigen at month 3. Pre-pandemic samples are used only as baseline. Medians are represented. *p<0.05, **p<0.01, ***p<0.001. IgG: Immunoglobulin G, RAU: relative antibody unit; WTS: whole trimeric spike; RBD: receptor-binding domain; S2: spike subunit 2; N: nucleocapsid; ME: membrane-envelop fusion protein.

### Age impacts the anti-SARS-CoV-2 and anti-HCoV IgG levels

As age is a known factor for COVID-19 symptoms severity, we then investigated the kinetics of SARS-CoV-2 specific IgG levels in relation to age groups. First, we compared the SARS-CoV-2 IgG levels from the 182 samples available at week 2 post-1^st^ PCR+ (**Fig 3A**). Interestingly, regardless of the specificity, the anti-SARS-CoV-2 IgG level in the pediatric and young adults’ population (< 25years) exhibited negative correlation with age while the adult population, i.e. 25-60years, exhibited positive correlation with age. The eldest population (> 60years) exhibited negative correlation with age. Overall, older age groups (44-58 and 60-95) mounted increased anti-SARS-CoV-2 antibody levels as compared to younger age groups (7-31 and 32-43, p <0.05, **Fig 3B** and **S3 Fig**). These observations were confirmed when analysing the maximum level of anti-SARS-CoV-2 between age groups (**Fig 3C** and **S4 Fig**): a significant gradual maximum level of anti-SARS-CoV-2 IgG was found across the different SARS-CoV-2 specific responses (p <0.01).

**Fig 3.**
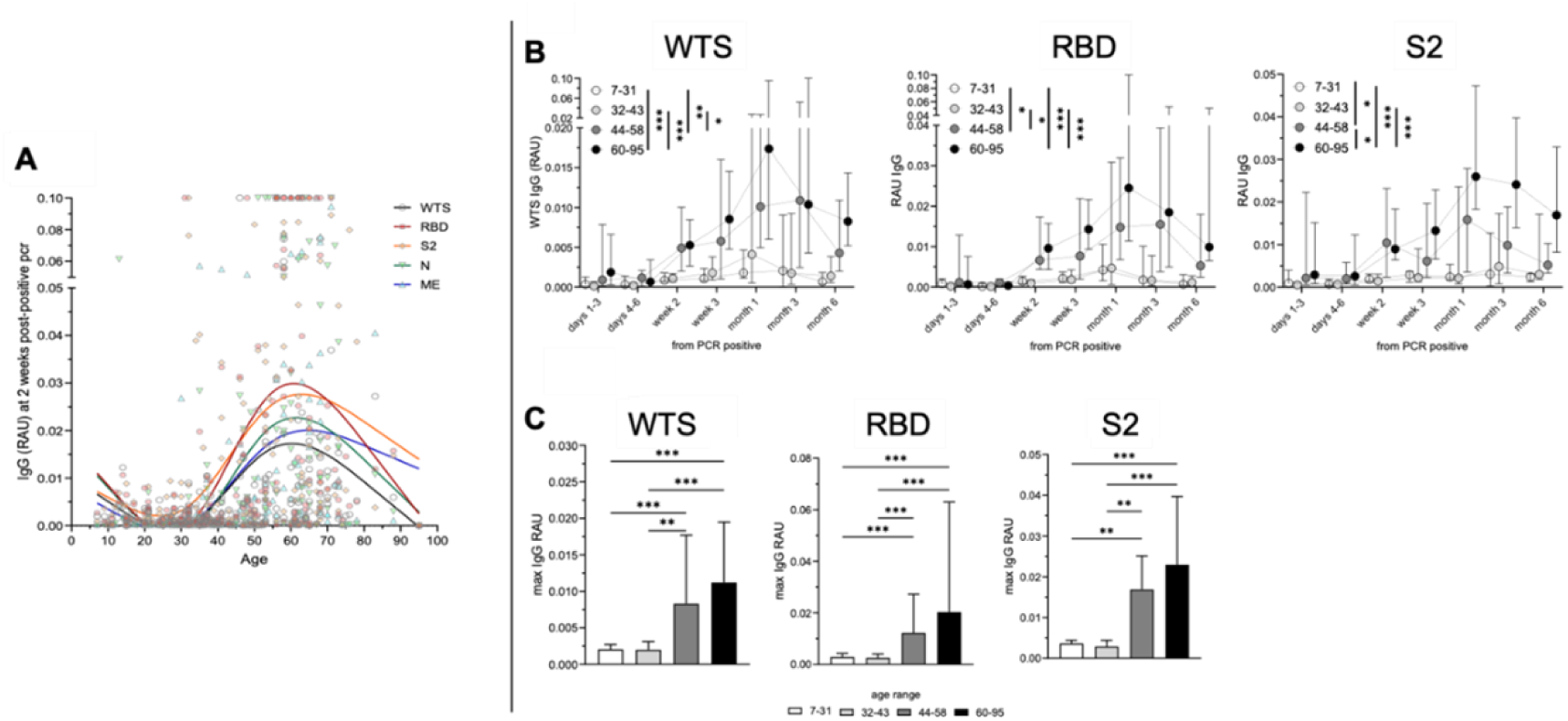
Correlation between age and SARS-CoV-2 IgG levels**. *A*,** Anti-SARS-CoV-2 IgG-based reactivity at 2 weeks post-PCR+ by age. Smoothing spline curves are represented for each IgG-specific reactivity. ***B***, Overtime reactivity against WTS, RBD and S2 antigens (RAU IgG). ***C***, Comparison of the peak level of anti-WTS, RBD, and S2 IgG (max IgG RAU) between age groups (quartile age range: 7–31, 32–43, 44–58, and 60–95 years). Kruskal-Wallis test with Dunn’s multiple comparison was used to compare peak levels between age groups. Overtime differences between groups were analyzed using two-way ANOVA with Šídák multiple comparison test. *p<0.05, **p<0.01, ***p<0.001. Medians with 95% CI are represented. IgG: Immunoglobulin G, RAU: relative antibody unit; WTS: whole trimeric spike; RBD: receptor-binding domain; S2: spike subunit 2.

The age-related HCoV IgG analysis showed that OC43 specific IgG levels pattern (**Fig 4A**) were similar to the SARS-CoV-2 one (**Fig 3A**) with negative correlation between anti-HCOV IgG levels and age in the youngest population (< 25years) and a positive correlation in the 25-60years adult population (**Fig 4A**). This pattern was not observed for the anti-HKU1 and anti-NL63 IgG levels. Accordingly, >44years groups had a significantly higher maximum anti-OC43 and anti-229E IgG level than younger groups (**Fig 4B**, p <0.01). No such observation was made regarding the anti-NL63 response and only a similar trend was observed in the anti-HKU1 IgG level between the youngest and oldest groups (p <0.05). Overtime post-1^st^ PCR+, anti-OC43, anti-229E (**Fig 4C**) as well anti-HKU1 IgG levels showed to be impacted by age (**S5 Fig**); especially the anti-229E response with the 60-95years group (p <0.001, **Fig 4C**). Note that in the oldest group the anti-OC43 IgG peaked earlier (week 2 post-PCR+) than the anti- WTS responses that peaked at month 1 post-PCR+ (**Fig 3B**).

**Fig 4.**
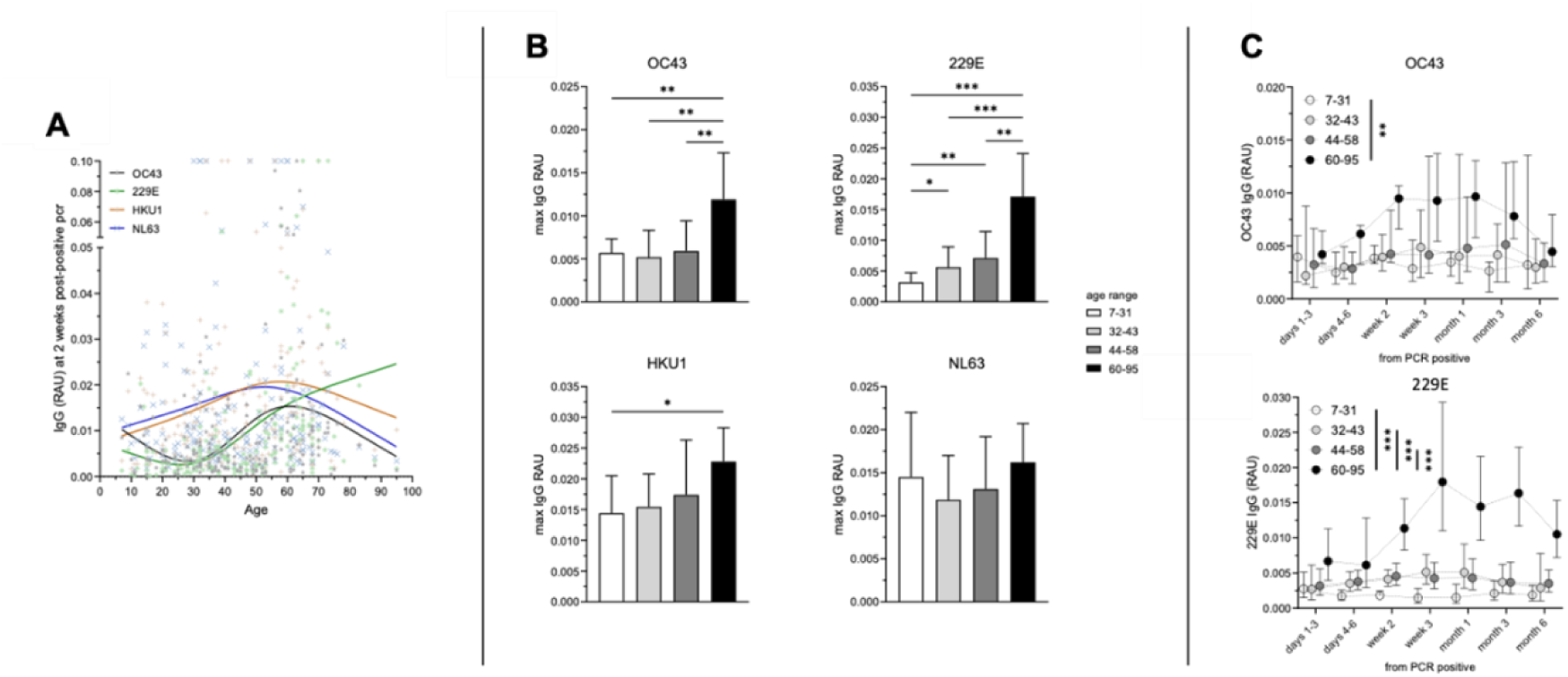
Correlation between age and HCoV IgG levels. ***A*,** Anti-HCoV IgG-based reactivity at 2 weeks post-PCR+ by age. Smoothing spline curves are represented for each IgG-specific reactivity. ***B***, Comparison of the peak level of anti-HCoV (OC43, 229E HKU1 and NL63) IgG (max IgG RAU) between age groups (quartile age range: 7–31, 32–43, 44– 58, and 60–95 years). ***C***, Overtime reactivity against OC43, and 229E antigens between the four age groups. Kruskal-Wallis test with Dunn’s multiple comparison was used to compare peak levels between age groups. Overtime differences between groups were analyzed using two-way ANOVA with Šídák multiple comparison test. *p<0.05, **p<0.01, ***p<0.001. Medians with 95% CI are represented. IgG: Immunoglobulin G, RAU: relative antibody unit.

### Kinetics of IgG-response among different clinical spectra of SARS-CoV-2

We further investigated the kinetics of SARS-CoV-2 specific IgG levels based on symptoms occurrence. First, we compared the highest recorded anti-SARS-CoV-2 antibody levels between symptomatic and asymptomatic patients (**Fig 5A**). Symptomatic patients showed higher maximum level of SARS-CoV- 2 specific IgG than asymptomatic patients against all SARS-CoV-2 proteins expect WTS. Additionally, while both groups showed an increased SARS-CoV-2 antibody level post-1^st^ PCR+, symptomatic patients mounted significantly higher SARS-CoV-2 specific IgG levels than asymptomatic patients for all SARS-CoV-2 proteins (**Fig 5B**, p<0.01 and **S6 Fig**). Regarding anti-HCoV responses, while symptomatic patients showed a higher maximum level of HCoV specific IgG than asymptomatic patients (**S7 Fig**); the anti-HCoV IgG levels kinetics did not differ between symptomatic and asymptomatic groups (**S7 Fig**). Since early SARS-CoV-2 specific IgG levels were similar (within the first week post-1^st^ PCR+) between symptomatic and asymptomatic groups, we compared the IgG kinetics between the different responses toward the specific anti-SARS-CoV-2 proteins. Within the asymptomatic group, no difference between the specific anti-SARS-CoV-2 IgG kinetics was observed (**Fig 5C**). In contrast, in the symptomatic group, kinetics of the specific antibody responses revealed that the anti-ME IgG levels were significantly lower than the other anti-SARS-CoV-2 response from month 1 post-1^st^ PCR+, especially compared to anti-RBD and anti-WTS IgG levels (**Fig 5C**, p <0.05).

**Fig 5.**
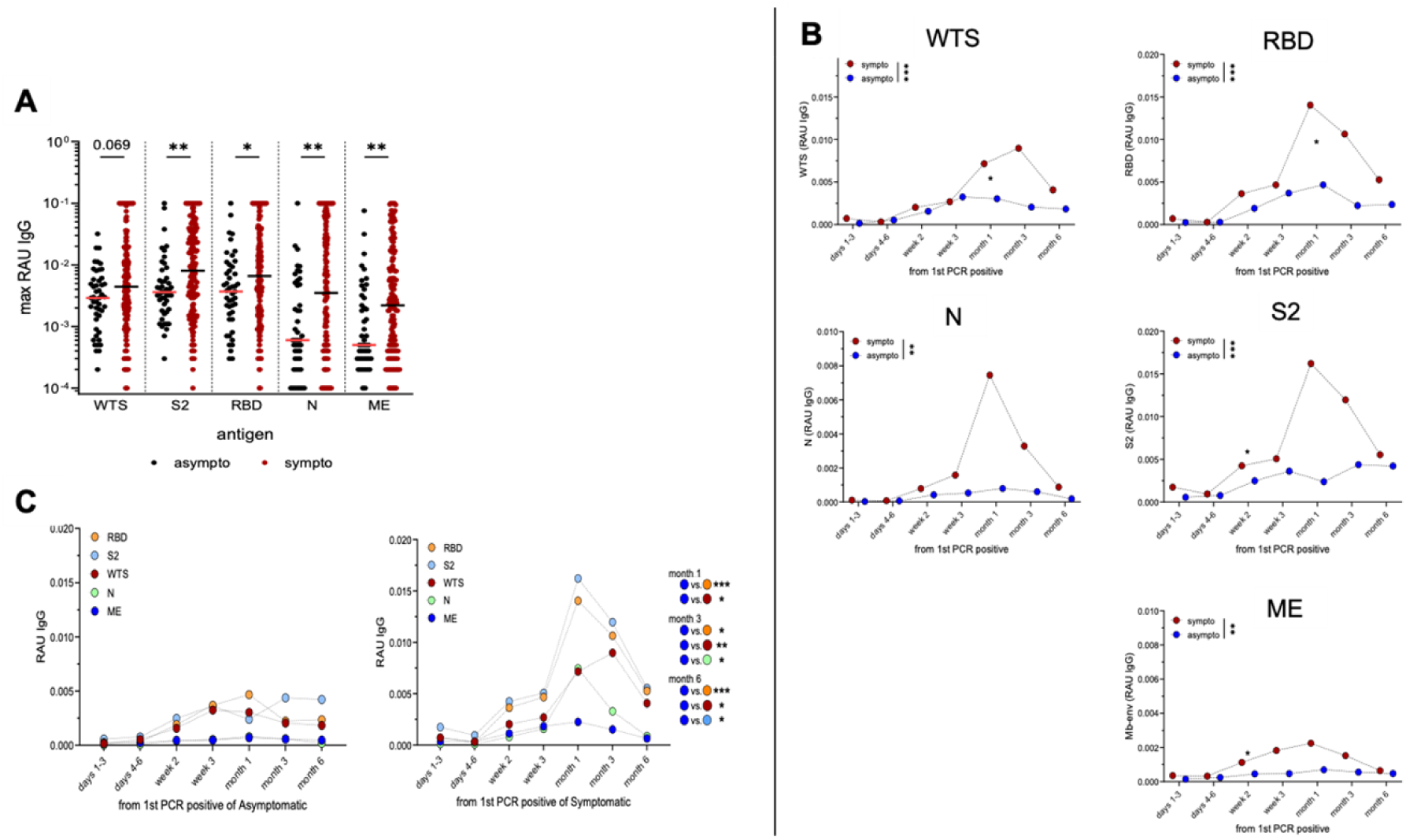
**Kinetics of SARS-CoV-2 IgG level from first positive PCR in correlation with symptoms**. ***A***, Comparison of the peak level of the 5 anti-SARS-CoV-2 IgG (max RAU IgG) for the asymptomatic (black, n=47) and symptomatic (red, n=157) groups. Medians are represented. Differences were analyzed using Mann–Whitney to detect differences between asymptomatic and symptomatic groups. ***B***, Overtime reactivity against SARS-CoV-2 antigens (RAU IgG) of asymptomatic (blue) and symptomatic (red) groups. **C**, Comparison kinetics between of the 5 anti-SARS-CoV-2 IgG in asymptomatic and symptomatic groups. Overtime differences between groups were analyzed using two-way ANOVA with Šídák multiple comparison test. *p<0.05, **p<0.01, ***p<0.001. IgG: Immunoglobulin G, RAU: relative antibody unit; WTS: whole trimeric spike; RBD: receptor-binding domain; S2: spike subunit 2; N: nucleocapsid; ME: membrane-envelop fusion protein.

### Correlation between anti-SARS-CoV-2 and anti-OC43 humoral responses in symptomatic patients

We then looked closely at the anti-HCoVs humoral kinetics (**Fig 6A**). Exclusively in symptomatic patients, OC43 IgG levels appeared to be significantly affected overtime post-1^st^ PCR+ (p <0.01) while no other anti-HCoVs humoral response changed post-infection. We also observed that the peak of IgG was different between antibody specificities (**Fig 6B**); indeed, in asymptomatic individuals, anti-WTS and anti-S2 responses peaked more rapidly than the other anti-SARS-CoV-2 responses (week 3 vs. month 1 post-1^st^ PCR+). Interestingly, in symptomatic patients, anti-OC43 showed the earliest peak at week 3, before any anti-SARS-CoV-2 peak that occurred at month 1 and later (**Fig 6B**). Correlation analysis between immune responses to SARS-CoV-2 and HCoVs 2 weeks post-1^st^ PCR revealed only a significant positive correlation between different anti-SARS-CoV-2 IgG responses in the asymptomatic cohort (**Fig 6C**). Interestingly, within the symptomatic group, on top of strong intra-SARS-CoV-2 IgG correlations, a notable positive correlation between anti-SARS-CoV-2 antibodies with anti-OC43 antibodies was highlighted, with the highest coefficient between OC43 (spike) and SARS-CoV-2 S2 (r=0.699, p <0.001, **Fig 6D**).

**Fig 6.**
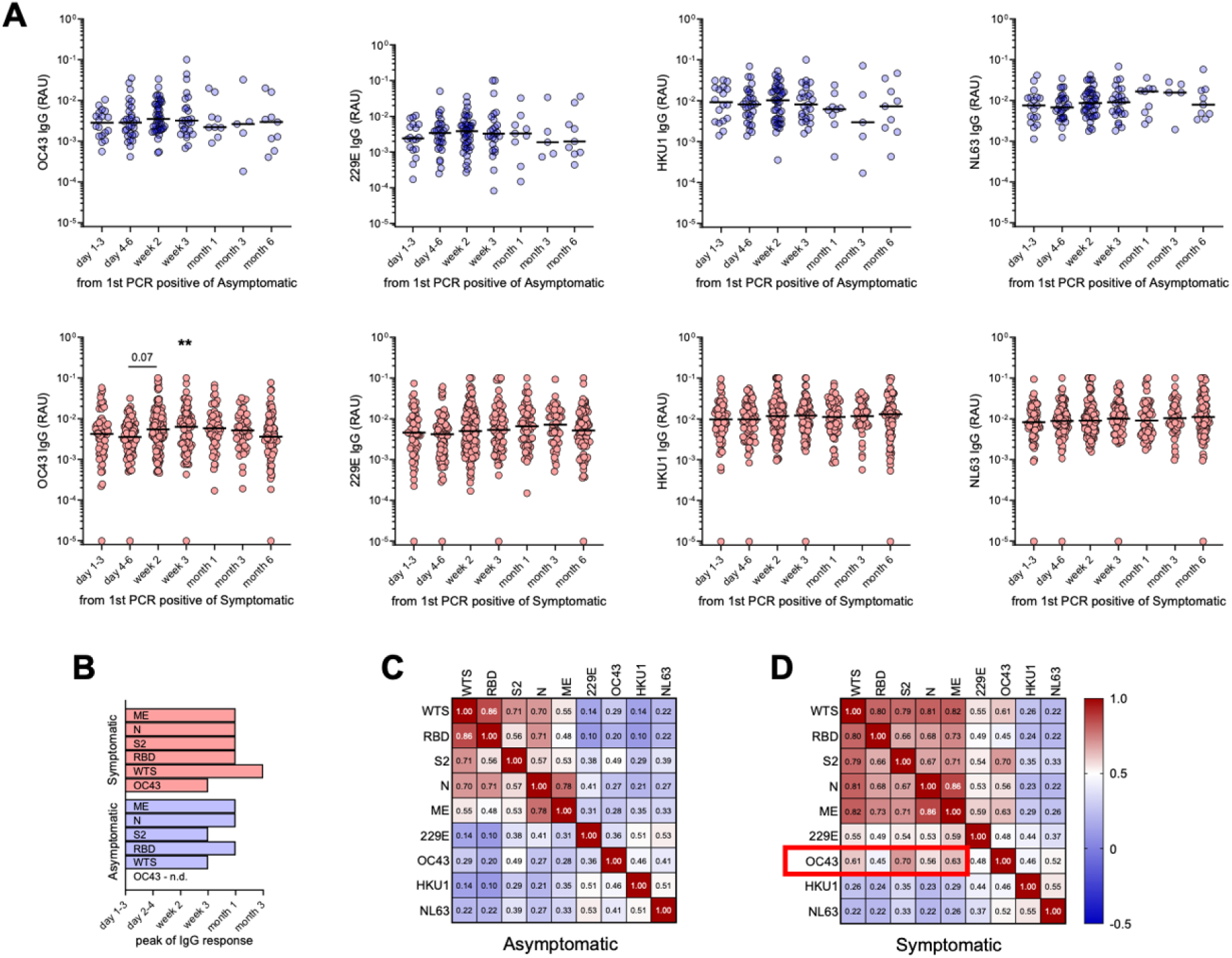
Correlation between anti-SARS-CoV-2 and anti-HCoV IgG levels in patients’ group at week 2 post-PCR+. ***A***, Kinetics of anti-HCoV IgG levels in asymptomatic and symptomatic group. Kruskal-Wallis test with Dunn’s multiple comparison was used to compare values of group of individual overtime. Medians are represented. ***B***, Time of specific IgG peaks post-PCR+ in asymptomatic and symptomatic groups. Heatmap for reactivity against 5 anti-SARS-CoV-2 and 4 anti-HCoV antigens in asymptomatic (***C***) and symptomatic patients (***D***). Correlation between two reactivity was assessed using Spearman correlation. For each correlation, “r” is indicated. IgG: Immunoglobulin G, RAU: relative antibody unit; WTS: whole trimeric spike; RBD: receptor-binding domain; S2: spike subunit 2; N: nucleocapsid; ME: membrane-envelop fusion protein.

## Discussion

We aimed to investigate potential cross-reactivity between the dynamics of SARS-CoV-2- and HCoV-specific IgG responses in a Senegalese population. As previously reported [9, 18], post-infection, anti-SARS-CoV-2 IgG levels rapidly increased and sustained up to six months, with peak levels at month one post-PCR+. While RBD and WTS showed stronger immunogenicity, the anti-N and anti-ME specific IgG waned more rapidly, excluding them for vaccine candidates.

In this Sub-Saharan cohort, the kinetics of the SARS-CoV-2 humoral immune response mirrored those from other continents regarding symptom severity and age. Kinetics of SARS-CoV-2-specific IgG levels were similar between symptomatic and asymptomatic groups, peaking at month one post-infection but the amplitude of response varied with the disease’s clinical spectrum, with asymptomatic patients showing lower antibody levels compared to symptomatic individuals [19, 20]. This could be due to shorter infection duration, a lower inflammatory response, and reduced antigen exposure [21] but also may indicate an alternative immune response, such as increased SARS-CoV-2-specific T cells [22]. The elderly population exhibited higher SARS-CoV-2 antibody levels than younger individuals, consistent with previous studies linking clinical outcomes of COVID-19 patients, age, and SARS-CoV-2 IgG levels early in the pandemic [23–25]. Nearly a quarter of our cohort was under 30, with 20 individuals below 20 years old, highlighting a trend toward higher antibody levels in this youngest group. To our knowledge, only 2 studies also described this reverse correlation between antibody response and age in a children-young adults’ cohort [25, 26]. This younger group may respond differently to SARS-CoV-2 infection due to more effective innate immunity and a respiratory tract preserved from pollution and underlying disorders [27].

We did not observe that HCoVs antibodies impacted the probability of SARS-CoV-2 infection, but we identified an increased anti-OC43 IgG response post-SARS-CoV-2 infection and an age-related antibody pattern similar between SARS-CoV-2 and the HCoVs OC43 and 229E. This suggests that a pre-existing anti-HCoV memory humoral immunity was triggered by the infection or serological cross-reactivity. A retrospective Senegalese epidemiological analysis (2012-2020) showed OC43 as the main circulating HCoVs (57.9% of tested) [16], supporting the possible existence of anti-OC43 memory immunity.

Symptomatic patients showed a rapid increase in anti-OC43 IgG, peaking earlier than any anti-SARS-CoV-2 humoral response post-infection. In contrast, asymptomatic patients did not exhibit an increased anti-OC43 humoral response but had an earlier anti-WTS and anti-S2 IgG peak. This earlier peak in symptomatic group supports the hypothesis of cross-reactivation of anti-OC43 immune memory. These differing kinetics between symptomatic and asymptomatic groups may indicate that (i) pre-existing anti- OC43 memory humoral immunity might be detrimental to efficient SARS-CoV-2 immunity in the Senegalese population, or (ii) the anti-OC43 IgG response remained invisible in asymptomatic group because of the overall weaker response. Cross-reactivity between OC43 and SARS-CoV-2 immune responses has been characterized at the cellular level, especially in CD4 T cells [28, 29]. However, there are inconsistent reports on OC43’s impact on SARS-CoV-2 pathology and immune response. Cellular immunity generated after OC43 infection showed immune reactivity against SARS-CoV-2 in young children (<6 years old) [30], which declined with age. Conversely, CY Lin et al. found that individuals with severe SARS-CoV-2 infections had more antibody-secreting B cells reactive to HCoV [31] and demonstrated -in a mouse model- that prior immunization with HCoV spike protein limited the anti-SARS-CoV-2 RBD response. This study aligns with our results, showing that SARS-CoV-2 infection induced an increase in anti-OC43. Higher levels of (OC43-) HCoV antibodies post-infection correlated with SARS-CoV-2 antibody levels and were associated with severity.

Several limitations should be taken into account. In our study, we couldn’t definitively describe how the SARS-CoV-2 humoral response is hindered by pre-existing anti-OC43 immunity. Two hypotheses are possible based on our observations: (i) the anti-OC43 response delays the anti-SARS-CoV-2 (especially WTS and S2) response, contributing to symptom occurrence; or (ii) the pre-existing anti-OC43 response boosts the overall immune response, including anti-SARS-CoV-2 humoral immunity, to a higher magnitude, leading to symptoms and possible disease severity. Additionally, the classification of patients was based on clinical presentation at inclusion, but the asymptomatic classification can vary as mild, or subclinical symptoms may have not been captured accurately. Furthermore, we also noticed a loss of samples overtime in the asymptomatic group, possibly due to the loss of interest from that group, that reduced the statistical power in the later timepoints of the study.

In conclusion, our study evaluated the kinetics of SARS-CoV-2 antibodies, showing that anti-SARS-CoV-2 IgG antibodies persist for at least six months post-infection, with a stronger humoral response observed in symptomatic patients. Age was also found to significantly influence antibody levels against both SARS-CoV-2 and certain HCoVs. The observed cross-reactivity between anti-SARS-CoV-2 and anti-OC43 antibodies, especially in symptomatic individuals, suggested a potential link between prior HCoV exposure and the clinical presentation of COVID-19, a field that warrants further investigation. These results could have important implications for future vaccination strategies and development including specific and cross-reactive epitopes [32] but also development of mathematical models to refine the prediction of SARS-CoV-2 variants transmission [33].

## Materials and methods

### Study Population and Design

Samples from SARS-CoV-2 RT-PCR-positive patients (n = 204) were obtained from COVID-19 patients enrolled during (i) a prospective case-ascertained study of COVID-19 suspected cases and their close contacts performed between March and October 2020 (n = 93), (ii) a multicentric non-interventional national cohort survey SEN-COV (n = 86) and (iii) a clinical therapeutic trial assessing the efficacy and safety of COVID-19 treatments in Senegal SEN-COV-FADJ (n = 25) (**S1 Text**). For each SARS-CoV-2 positive patients, socio-demographic (age and sex), epidemiological (sampling date, onset of symptoms date), clinical data, symptoms and comorbidities were collected (**S1 Table**). Based on their clinical status at inclusion, SARS-CoV-2 positive patients were classified as asymptomatic (n = 47) or symptomatic (n = 157). Sex and age matched pre-SARS-COV-2 emergence plasma samples were also obtained from a longitudinal cohort survey conducted in Dielmo (Fatick region, Toubacouta district, Senegal), and ongoing since 1990 [34]. The samples used in this study were collected during a cross-sectional survey performed in June 2018.

### Ethics Statement

For the longitudinal cohort survey conducted in Dielmo, the retrospective use of data and serum samples for immunological analysis in the context of COVID-19 pandemic was approved by the Senegalese National Ethics Committee for Research in Health (reference number 00000007/MSAS/CNERS/Sec 26 January 2021). Individual written consents were obtained from the Dielmo villagers for this purpose. As part of the national COVID-19 pandemic surveillance in Senegal, approval by the National Ethics Committee of the Ministry of Health was not required. Data and samples were collected for surveillance purposes and were anonymized. However, oral informed consent was systematically obtained for the retrospective use of their data and samples for surveillance and/or research purposes. For the SEN-COV and SEN-COV-FADJ cohort survey, ethical approval was obtained from the Senegalese National Ethics Committee for Research in Health (CNERS, reference number 00000068/MSAS/CNERS/Sec, 10 April 2020) [35]. All patients included in these two cohorts provided written informed consent. This study was conducted following good clinical practice and according to the Declaration of Helsinki.

### Sample Collection

Naso/oropharyngeal swabs and blood samples were collected prospectively at inclusion and during the hospitalization period. Blood samples were also collected during follow-up of patients up to 6 months after their enrollment. Nasopharyngeal samples were used for SARS-CoV-2 Real Time PCR detection. A total number of 705 blood samples were collected over a period of up to 6 months from the first positive PCR. Plasma sample collection was performed between March 2020 and January 2022 (**Fig 1A**). Patients enrolled after November 2020 (n = 39) were excluded due to vaccination initiation and start of the 2^nd^ wave of SARS-CoV-2 (Beta) in Senegal (**Fig 1B**). Blood samples were collected in a 4 ml vacutainer EDTA tubes at inclusion (day 0-1 post-PCR+) and every 48-72 hours until hospital discharge and then at 3 timepoints (month 1, month 3, and month 6 post-PCR+) when possible. For some patients (n =20, mostly contact individuals or suspected cases), plasma samples collected before SARS-CoV-2 infection confirmation by positive naso/oropharyngeal RT-PCR test, were included. Plasma samples were obtained after blood centrifugation at 2500 rpm for 5 min and stored at -20°C until use. Data were recorded on standardized investigation forms and sent to Institut Pasteur de Dakar.

### IgG response analysis by bead-based multiplex assay

The bead-based multiplex serological assay use in this study includes five SARS-CoV-2 antigens of the ancestral variant: whole trimeric spike (WTS), spike’s receptor-binding domain (RBD), and spike subunit 2 (S2), nucleocapsid protein (N), and a membrane-envelope fusion protein (mb-env); and the four HCoV (spike proteins of NL63, 229E, HKU1, OC43). The assay was conducted as described previously and in **S1 Text** [36, 37].

### Statistical Analysis

Data analysis was performed with GraphPad Prism software (version 9). For statistical purposes, patients were then divided by age quartiles (**Fig 1B**). The frequency of symptomatic patients within each age group (quartiles) was compared through the Chi-squared test. Two-dimensional hierarchical clustering was performed using Online CIMminer using the Euclidian distance method and the average linkage cluster algorithm. Differences between 2 or more unpaired groups were analysed using Mann– Whitney or Kruskal-Wallis test with Dunn’s multiple comparison. Two-way ANOVA with Šídák multiple comparison test was used to compare the different kinetics between groups of patients (age or symptoms). Correlation between IgG-based reactivity was assessed by non-parametric Spearman correlation “r”. Correlation between age and antigen specific IgG level was represented using spline curve with smoothing spline of 4 knots. The threshold of significance was assessed at 0.05. medians are represented.

## Acknowledgments

The authors would like to express their deep gratitude to the clinical research teams of the Infectious and Tropical Diseases Department at Fann Hospital, Hôpital Principal de Dakar, Dalal Diam Hospital in Guédiawaye (including physicians and clinical investigators, site managers, regulatory staff, clinical research coordinators), and the Ministry of Health and Social Action of Senegal for their close collaboration with the Institut Pasteur de Dakar, their invaluable assistance in recruiting participants, and their logistical support throughout this study. We also extend our sincere thanks to the study participants for their vital contribution and to the laboratory personnel, without which this research would not have been possible.

## Supporting information legends

**S1 Fig**. **Hierarchical clustering of IgG reactivity toward SARS-CoV-2 and HCoV antigens in 20 pre-PCR+ samples** (y axis: asymptomatic A, n=11; symptomatic S, n=9). Medians are represented. Spike WT: whole trimeric spike; RBD: receptor-binding domain; S2: subunit 2; N: nucleocapsid; ME: membrane-envelop

**S2 Fig. Overall overtime relative IgG-based reactivity toward SARS-CoV-2 and HCoV antigens from 1^st^ positive PCR**. Medians are represented. Spike WT: whole trimeric spike; RBD: receptor-binding domain; S2: subunit 2; N: nucleocapsid; ME: membrane-envelop.

**S3 Fig**. **Overtime reactivity against S2, RBD, N and ME antigens (RAU IgG) between 4 age groups (quartile age range 7-31, 32-43, 44-58 and 60-95).** Overtime differences between groups were analysed using two-way ANOVA with Šídák multiple comparison test. Kruskal-Wallis test with Dunn’s multiple comparison was used to compare between peak levels between age groups. *p<0.05, **p<0.01, ***p<0.001. Medians with 95% CI are represented. IgG: Immunoglobulin G. RAU: relative antibody unit; RBD: receptor-binding domain; S2: subunit 2; N: nucleocapsid; ME: membrane-envelop.

**S4 Fig. Comparison of the available peak level of anti-N and mb-env IgG (max RAU IgG) between age groups.** IgG: Immunoglobulin G. RAU: relative antibody unit; RBD: receptor-binding domain; S2: subunit 2; N: nucleocapsid; ME: membrane-envelop.

**S5 Fig. Overtime reactivity against HKU1 and NL63 antigens (RAU IgG) between 4 age groups (quartile age range 7-31, 32-43, 44-58 and 60-95).** Medians with 95% CI are represented and overtime differences between groups were analysed using two-way ANOVA with Šídák multiple comparison test. *p<0.05, **p<0.01. IgG: Immunoglobulin G. RAU: relative antibody unit.

**S6 Fig. Kinetics of anti-SARS-CoV-2 IgG levels in asymptomatic individuals.** Kruskal-Wallis test with Dunn’s multiple comparison was used to compare values of group of individuals overtime. Data analysed were performed from the peak of the response. Overtime differences between groups were analysed using two-way ANOVA with Šídák multiple comparison test. *p<0.05, **p<0.01. IgG: Immunoglobulin G. RAU: relative antibody unit.

**S7 Fig. Comparison of the available peak level of the 4 anti-HCoV (max RAU IgG) for the asymptomatic (black, n=47) and symptomatic (red, n=157) groups.** Differences were analysed using Mann–Whitney to detect differences between asymptomatic and symptomatic groups. Overtime reactivity against HCoV antigens (RAU IgG) of asymptomatic (blue) and symptomatic (red) groups (C). Medians are represented. Overtime differences between groups were analysed using two-way ANOVA with Šídák multiple comparison test. *p<0.05, **p<0.01. IgG: Immunoglobulin G. RAU: relative antibody unit.

